# MAFB is essential for the maintenance of adult human α-cell identity and glucagon secretion

**DOI:** 10.64898/2026.08.08.743687

**Authors:** KC Coate, J Liu, M Guo, X Tong, VMN Coykendall, C Harmelink, N Dey, G Reynolds, N Mohanty, RE Jenkins, R Aramandla, JP Cartailler, AC Powers, PE MacDonald, SK Kim, RW Stein

**Affiliations:** Department of Molecular Physiology and Biophysics, Vanderbilt University School of Medicine, Nashville, TN; Department of Medicine, Division of Diabetes, Endocrinology and Metabolism, Vanderbilt University Medical Center, Nashville, TN; Department of Veterans Affairs, Tennessee Valley Healthcare System, Nashville, TN; Creative Data Solutions, Vanderbilt Center for Stem Cell Biology, Vanderbilt University, Nashville, TN, 37232, USA; Department of Pharmacology and Alberta Diabetes Institute, University of Alberta, Edmonton, Alberta, Canada; Department of Developmental Biology, Stanford University School of Medicine, Stanford, CA; Stanford Diabetes Research Center, Stanford University School of Medicine, Stanford, CA; Department of Medicine, Stanford University School of Medicine, Stanford, CA; Department of Pediatrics, Stanford University School of Medicine, Stanford, CA

## Abstract

Dysregulated hormone secretion and erosion of endocrine cell identity are features of type 1 and type 2 diabetes, but the transcriptional programs maintaining adult human islet identity and function remain poorly defined. The large MAF transcription factor MAFB is expressed in human α- and β-cells, marks their most functionally mature subpopulations, and is downregulated in diabetes, but its role in adult human islets has not been tested directly. Using shRNA-mediated MAFB knockdown (KD) in whole and CD26^+^ α-cell-enriched human pseudoislets, we found that whole pseudoislet MAFB KD impaired glucagon synthesis and secretion while only modestly reducing insulin content and cAMP-potentiated insulin release. Single-cell profiling detected no β-cell transcriptional response beyond MAFB KD itself, consistent with buffering by the related β-cell-enriched MAFA transcription factor. In contrast, α-cell-restricted MAFB KD unmasked a cell-autonomous requirement for MAFB in stimulus-secretion coupling. MAFB deficiency also destabilized α-cell identity, downregulating canonical α-cell and neuroendocrine secretory genes while ectopically inducing mesenchymal and extracellular matrix remodeling programs. In addition, MAFB-dependent downregulation of electron transport chain genes was confined to a large α-cell subcluster, manifesting as impaired islet-wide mitochondrial respiration within the broader α-cell population. Together, these findings identify MAFB as an essential adult human α-cell maintenance factor that links diabetes-associated downregulation to impaired glucagon secretion, α-cell identity erosion, and mitochondrial dysfunction.

**RESEARCH IN CONTEXT:**

- **What is already known about this subject?**

- MAFB is expressed in adult human α- and β-cells, marks their most functionally mature subpopulations, and is downregulated in type 1 and type 2 diabetes
- In human stem cell models, MAFB is essential for generating insulin-producing β-like cells, whereas glucagon-producing α-like cells are reduced but still formed
- Neither model addresses adult human islets: rodent MafB becomes α-cell restricted after birth, and stem cell models capture differentiation, not maintenance
- **What is the key question?**

- Is MAFB required to maintain identity and secretory function in adult human islet cells?
- **What are the new findings?**

- MAFB knockdown in primary human pseudoislets impaired glucagon synthesis and secretion but minimally affected β-cells, consistent with buffering by MAFA
- Knockdown in CD26+ α-cell-enriched pseudoislets revealed a cell-autonomous requirement for MAFB in stimulus-secretion coupling, and destabilized α-cell identity by inducing mesenchymal and extracellular matrix programs
- MAFB loss downregulated electron transport chain genes in the largest α-cell subcluster and reduced mitochondrial respiration
- **How might this impact on clinical practice in the foreseeable future?**

- Preserving MAFB activity in adult human α-cells may represent a strategy to limit α-cell dysfunction in diabetes

## INTRODUCTION

MAFB (V-Maf avian musculoaponeurotic fibrosarcoma oncogene homolog B) is a basic leucine zipper transcription factor (TF) of the large MAF subfamily that is expressed across a broad set of tissues, including macrophages, podocytes, osteoclasts, lens epithelium, the developing hindbrain, and pancreatic islets.^1^ In humans, heterozygous MAFB variants cause multicentric carpotarsal osteolysis syndrome, underscoring its broader developmental and tissue-specific functions.^2,3^ In the pancreatic islet, MAFB is downregulated in α- and β-cells in both type 1 and type 2 diabetes (T1D and T2D), and is among the earliest islet TFs to be downregulated in response to cellular stress.^4–8^ These observations highlight context-dependent roles for MAFB across diverse cell types and suggest that loss may contribute directly to endocrine cell dysfunction in diabetes.

Within the pancreatic islet, MAFB is among the last TFs activated during endocrine cell maturation and is enriched in the most prevalent and functionally mature α- and β-cell subpopulations.^9,10^ MAFB transcripts have also been detected in adult human δ-cells^11,12^ by single cell RNA-sequencing (scRNA-seq), but its role in this relatively minor islet subpopulation remains uncharacterized. Although many islet-enriched TFs have been extensively studied in rodent models of embryonic endocrine cell development and function, MAFB is unique because its expression pattern differs between rodents and humans: rodent MafB is expressed in both α-and β-cells during development but becomes restricted to α-cells postnatally, whereas human MAFB is expressed in both α- and β-cells during development and throughout adulthood.^9,10,13–17^ Rodent models therefore cannot address MAFB function in adult human β-cells. In human embryonic stem cells (hESC), MAFB deletion spares early pancreatic and endocrine progenitor specification but redirects endocrine allocation toward somatostatin- and pancreatic polypeptide-producing cells, with insulin-producing β-like cells nearly abolished and non-functional after transplantation, whereas glucagon-producing α-like cells are reduced approximately 2-fold yet retain glucose-regulated secretion.^10^ Because MAFB expression persists in adult human α- and β-cells, it likely serves roles in mature islet cells that neither system can reveal. Interpreting that role is further complicated by potential redundancy within the large MAF family: MAFB shares cis-element target sequences with MAFA, which is highly enriched in β-cells but essentially absent from other islet cell types.^16,18^ Whether adult human β-cells depend directly on MAFB or are compensated by MAFA remains an open question.

Here, we used a human pseudoislet approach^19,20^ to define the regulatory role of MAFB in adult human α- and β-cells. Notably, MAFB knockdown (KD) in whole pseudoislets produced minimal impairment in insulin secretion and the β-cell transcriptome. In contrast, MAFB KD in α-cells disrupted glucagon expression and secretion, markedly destabilized α-cell molecular identity, and reorganized α-cell subpopulation-specific gene programs. Together, these findings establish MAFB as an essential cell-autonomous regulator of adult human α-cells and provide a framework linking diabetes-associated MAFB downregulation to α-cell dysfunction.

## METHODS

### Human islet procurement and culture

Deidentified human pancreatic islets (n = 25 preparations, Supp. Table 1) from previously healthy donors were procured through the Integrated Islet Distribution Program (IIDP; https://iidp.coh.org/), Alberta Diabetes Institute IsletCore (https://www.bcell.org/adi-isletcore.html)^21^, Human Pancreas Analysis Program (HPAP)^22–25^ (https://hpap.pmacs.upenn.edu/), or isolated at the Institute of Cellular Therapeutics of the Allegheny Health Network. Primary human islets were hand-picked to purity and cultured in CMRL 1066 media (5.5 mM glucose, 10% FBS, 1% penicillin/streptomycin, 2 mM L-glutamine) in 5% CO_2_ at 37°C for less than 24 hours before pseudoislet generation.

### Lentivirus production

Lentiviral constructs coexpressing a fluorescent transgene and short hairpin RNA (shRNA) directed against human MAFB (MAFB shRNA; GCCCAGTCTTGCAGGTATAAAC, VB180215-1122jqv) or a scrambled nontargeting control sequence (SCR shRNA; CCTAAGGTTAAGTCGCCCTCG, VB190410-1184rda) were obtained from Vector Builder. Lentiviruses were produced as described previously17 by transfection of HEK 293T cells, and supernatants were collected and concentrated using PEG-it Virus Precipitation Solution (System Biosciences). Pellets were resuspended in DMEM, aliquoted, and stored at −80°C prior to transduction. Viral particle amount was quantified with the Lenti-X p24 Rapid Titer Kit (Takara Bio, Mountain View, CA).

### Human pseudoislet formation: whole islet

Human pseudoislets were generated as previously described.^19^ Briefly, intact human islets were dispersed into single cells by enzymatic digestion with 0.025% HyClone trypsin (Thermo Scientific), counted with an automated Countess II cell counter, and transduced with lentivirus at a multiplicity of infection of 500 in Vanderbilt Pseudoislet Medium (VPM), and seeded into wells at 2000 cells per 200 µl in CellCarrier Spheroid ULA 96-well Microplates (Revvity). Pseudoislets were allowed to reaggregate for six days prior to analysis. To assess *MAFB* knockdown, RNA was extracted from pseudoislets using an RNAqueous™-Micro Kit (Invitrogen). cDNA was generated using the iScript cDNA synthesis kit (Bio-Rad) and quantitative RT-PCR (qRT-PCR) assays performed using the LightCycler FastStart DNA Master PLUS SYBR Green kit (Roche) and a LightCycler PCR instrument (Roche). qRT-PCR primers are listed in Supp. Table 2. qRT-PCR results were normalized to *ACTB* and scramble control samples using the comparative ΔΔCt method.

### Human pseudoislet formation: α-cell-enriched

Dispersed human islet cells were labeled with a PE-conjugated anti-human CD26 antibody (BioLegend) and separated using the EasySep Release Human PE Positive Selection Kit (STEMCELL Technologies) following the manufacturer’s protocol as previously described.^26^ CD26⁺ cells were transduced with MAFB KD (αMAFB^KD^) or SCR (αSCR) control lentiviruses (500 MOI), seeded into wells at 2000 cells per 200 µl in CellCarrier Spheroid ULA 96-well Microplates (Revvity), and allowed to reaggregate for 6 days before analyses. Enrichment of GCG-expressing cells was verified by qRT-PCR comparing *GCG*, *INS*, and *SST* mRNA levels in CD26⁺ and CD26⁻ fractions.

### Static incubation hormone secretion assays

Pseudoislets (16-20 islet equivalents [IEQ]/well) were transferred to cell culture inserts (12.0 µm, 12 mm diameter; Millicell) in a 24-well plate and equilibrated for at least 30 min in 1.0 mL of DMEM (D5030; Sigma) media containing 4.7 mM HEPES, 38.1 mM sodium bicarbonate, 4.0 mM L-glutamine, 1.0 mM sodium pyruvate, and 0.1% BSA (heat shock fraction, suitable for RIA) at 5.6 mM glucose.^27^ These were then transferred to DMEM media containing the secretagogue of interest (i.e., 5.6 mM glucose [basal], 1.7 mM glucose, 16.7 mM glucose, 16.7 mM glucose + 100 μM isobutylmethylxanthine [IBMX], and 1.7 mM glucose + 20 mM L-arginine) for 1 hour (3 technical replicates per condition per donor). Supernatants were collected and pseudoislets were lysed in 200 µl of acid alcohol (5.5 mL 95% ethanol, 50 µl 12N HCl) for islet hormone extraction.^27^ Glucagon and insulin were measured by ELISA (Mercodia), and hormone secretion normalized to pseudoislet volume (IEQ) or to total hormone content (fractional secretion).

### Immunofluorescence and cell composition analysis

Immunohistochemical analysis was performed on 8-µm cryosections of pseudoislets embedded in collagen I gels, as previously described.^28–30^ Briefly, pseudoislets were immobilized in collagen gels and fixed in freshly prepared 4% paraformaldehyde (PFA) in 10 mM PBS for 15 min at 37°C, followed by 45 min on ice. Samples were then washed three times in cold 10 mM PBS over 1 hour, equilibrated in 30% sucrose/10 mM PBS overnight at 4°C, and cryopreserved in optimum cutting temperature compound at −80°C. For immunostaining, cryosections were permeabilized in 0.2% Triton X-100 for 15 min at room temperature, blocked in 5% normal donkey serum for 1.5 hours, and incubated with primary antibodies overnight at 4°C. Secondary antibodies were subsequently applied for 1 hour at room temperature. All antibodies (Supp. Table 3) were diluted in 10 mM PBS containing 1% BSA and 0.1% Triton X-100. Sections were mounted with AquaPoly/Mount, and digital images were acquired on an Olympus FV3000 confocal laser scanning microscope at 20x with up to 3x digital zoom. Images were processed using HighPlex FL v3.2.1 within HALO software (Indica Labs).

### Transmission electron microscopy

Whole pseudoislets were fixed in freshly prepared 2% glutaraldehyde and 2% PFA in 0.1 M sodium cacodylate buffer overnight at 4°C, washed in 10 mM PBS, and sequentially post-fixed in 1% tannic acid, 1% OsO4, and stained with 1% uranyl acetate for 1 hour each. Samples were dehydrated in a graded ethanol series and infiltrated with Epon-812 using propylene oxide as a transition solvent. The resin was polymerized at 60°C for 48 hours and ultrathin sections were cut at 70 nm nominal thickness using a Leica ultramicrotome. Samples were imaged on a JEOL 2100 Plus transmission electron microscope operated at 200 KeV equipped with an AMT Nanosprint 15 MKII CMOS camera. Pseudoislet α-cells were identified by their characteristic glucagon granule morphology.^31^ Glucagon granule number and fractional granule area per α-cell were quantified from transmission electron micrographs using pixel classifier-based segmentation in the Labkit plugin^32^ for Fiji/Image J. For each α-cell, cytoplasmic area was defined as whole cell area minus nuclear area, and fractional granule area was calculated as total granule area divided by cytoplasmic area. Up to 10 α-cells were analyzed per condition per donor across 3 independent donors, yielding 27 scramble shRNA α-cells (8710 glucagon granules) and 31 MAFB shRNA α-cells (7082 glucagon granules) for analysis.

### Bulk RNA-sequencing, data processing, and analysis

RNA was extracted from CD26^+^ αMAFB^KD^ and αSCR control samples from 7 adult human donors without diabetes (Supp. Table 1) using an RNAqueous™-Micro Kit (Invitrogen). Quality control analysis was performed on all RNA samples using an Agilent 2100 Bioanalyzer. Samples with an RNA Integrity Number (RIN) ≥ 7.0 (mean ± SEM: 9.1 ± 0.4) were used for library preparation. mRNA enrichment and cDNA libraries were prepared using stranded mRNA (polyA-selected) library preparation kit (NEB). Libraries were sequenced as paired-end 150-bp reads on the Illumina NovaSeq 6000 targeting an average depth of 50 million reads per sample. RNA sequencing reads were adapter-trimmed and quality-filtered using Trimgalore v0.6.7.^33^ An alignment reference was generated from the GRCh38 human genome and GENCODE comprehensive gene annotations (Release 26), to which trimmed reads were aligned and counted using Spliced Transcripts Alignment to a Reference (STAR) v2.7.9a^34^ with the quantMode geneCounts parameter. On average, 18 million uniquely mapped reads were acquired per sample. DESeq2 package v1.36.0^35^ was used to perform sample-level quality control, low count filtering, normalization and downstream differential gene expression analysis. The paired sample analysis, comprising matched αMAFB^KD^ and αSCR conditions by donor, was handled via the design formula “∼ donor + condition”, which resulted in 70% PC1 variance with clear separation between conditions. Genomic features counted fewer than five times across at least three samples were removed. False discovery rate adjusted for multiple hypothesis testing with Benjamini-Hochberg procedure *p* value < 0.05 and log2 fold change > 0.5 was used to define differentially expressed genes (DEGs). Gene set overlap analysis was performed using the Human MSigDB database v2024^36,37^ ‘Compute Overlaps’ tool. The top 200 most significant up- or down-regulated genes from CD26^+^ αMAFB^KD^ versus αSCR control samples were queried against all MSigDB collections. Overlap significance was determined using hypergeometric distribution testing with Benjamini-Hochberg FDR correction (q < 0.05).

### Fixation of single cells for chromium fixed RNA profiling

On day 6 post-transduction, scramble shRNA control and MAFB KD whole pseudoislets (150 per condition) prepared from two adult donors without diabetes (Supp. Table 1) were dispersed into single cells and fixed according to the manufacturers’ protocol (Fixation of Cells & Nuclei for Chromium Fixed RNA Profiling, CG000478, Rev D, https://www.10xgenomics.com/). Briefly, pseudoislets were washed twice with sterile DPBS containing 2 mM EDTA, dispersed in freshly prepared 0.025% Hyclone trypsin (250 µl per sample) for 7-10 minutes at room temperature by gentle trituration, quenched with 1.75 mL CMRL 1066 media (recipe above), and washed once with sterile DPBS. The supernatant was removed leaving ∼30 µl, and 1 ml of freshly-prepared fixation buffer (791.9 µl nuclease-free water + 100 µl conc. Fix & Perm Buffer [10x genomics PN-2000517] + 108.1 µl 37% formaldehyde), 0.1 volume of pre-warmed Enhancer (10x Genomics PN-2000482), and 50% glycerol (final concentration of 10%) was added to each sample, pipetted five times to mix, stored at 4°C for 20h and −80°C for ∼30 days.

### Single-cell RNA-sequencing, data processing, and analysis

Single-cell RNA-seq (scRNA-seq) libraries were prepared using the 10X Genomics Fixed RNA Profiling (Gene Expression FLEX) chemistry and sequenced on the Illumina NovaSeq 6000 platform targeting 20,000 reads per cell for the gene expression libraries. Raw sequencing reads were aligned and demultiplexed using Cell Ranger Cloud (10x Genomics; https://www.10xgenomics.com/), with probe-based cell calling and doublet removal based on Flex kit-specific oligo-tagged probes. Cell-containing droplets were identified using Dropkick^38^, and ambient RNA was corrected using CellBender.^39^ The resulting count matrices were loaded into Seurat^40^ (v4.4.0) in R. Starting from 16,291 genes across 8,882 cells, quality control filtering removed cells with mitochondrial content >10%, fewer than 250 or more than 10,000 detected features, and fewer than 1,000 or more than 100,000 unique molecular identifiers (UMIs), retaining 8,319 cells. Doublets were identified and removed using scDblFinder^41^ with cluster-based automatic doublet rate estimation, yielding 7,714 single cells for downstream analysis. Gene expression counts were log-normalized (scale factor = 10,000) and the top 2,000 variable features were selected via variance-stabilizing transformation. Samples were integrated using Harmony^42^ on sample identity across 30 principal components and then Uniform Manifold Approximation and Projection (UMAP) was computed on the Harmony-corrected embeddings. Graph-based clustering at resolution 0.5 identified 16 clusters, which were manually annotated into 9 cell populations based on canonical marker gene expression: α-, β-, δ/PP/ε cells, acinar cells, ductal cells, stellate cells, Schwann cells, mast cells, and immune/endothelial cells. Islet α-cells constituted most of the dataset (5,059 cells), followed by β-cells (1,491 cells). For differential gene expression analysis of α-cells, an α-cell subset was re-clustered using Harmony^42^ integration at resolution 0.2, yielding 6 subclusters. Proliferating and minor clusters were excluded, leaving 4,464 α-cells across 5 subclusters for differential expression (DE) testing. Single-cell DE between MAFB KD and control was performed using MAST^43^ (minimum detection rate = 20%), identifying 238 differentially expressed genes (adjusted p < 0.05; 49 downregulated, 189 upregulated). Per-subcluster DE within α-cell subpopulations C0 – C3 was also conducted using both Wilcoxon rank-sum and MAST^43^ tests at the single-cell level. Functional enrichment analysis of C0 – C3 was performed using g:Profiler.^44^ α-Cell subcluster C4 was excluded from these analyses because no genes met the adjusted p < 0.05 threshold for differential expression (subcluster C4 contained only 33 cells). This work also used data acquired from the Human Pancreas Analysis Program (HPAP; (https://hpap.pmacs.upenn.edu/) (RRID:SCR_016202), supported by the Human Islet Research Network (HIRN; RRID:SCR_014393)

### Seahorse XF pro pseudoislet respirometry

Human pseudoislets were generated from six adult donors without diabetes (Supp. Table 1; CD26^+^ αMAFB^KD^ and αSCR control pseudoislets, n=4; MAFB KD and SCR shRNA whole pseudoislets, n=2). Pseudoislets were seeded into XF96 spheroid plates (Agilent) as previously described^45^ in assay medium consisting of DMEM (D5030; Sigma) supplemented with 4.7 mM HEPES, 4.0 mM L-glutamine, 1.0 mM sodium pyruvate, 0.02% BSA, and 3.0 mM glucose. Prior to the assay, plates were equilibrated for 1 hour at 37°C in a non-CO₂ incubator. Mitochondrial respiration was assessed using the Seahorse XF Pro extracellular flux analyzer and the Mito Stress Test kit (Agilent). Basal oxygen consumption rate (OCR) was measured at 3 mM glucose, after which pseudoislets were sequentially challenged with oligomycin (4.5 μM), FCCP (2.0 μM), and rotenone/antimycin A (2.5 μM). OCR values were normalized to islet volume (IEQ). Bioenergetics parameters were calculated as follows. Non-mitochondrial respiration was defined as the minimum OCR value after rotenone/antimycin A injection. Basal respiration was calculated as the last OCR value before oligomycin injection minus non-mitochondrial respiration, and maximal respiration was calculated as the maximum OCR value after FCCP injection minus non-mitochondrial respiration. ATP-production coupled respiration was calculated as the last OCR value before oligomycin injection minus the minimum OCR value after oligomycin injection.

### Study approval

All studies were conducted with deidentified human pancreatic islets and are therefore not classified as human subject research by the Vanderbilt University Institutional Review Board.

### Statistical analyses and data visualization

Data are presented as mean ± SEM unless otherwise specified. Statistical comparisons between groups were performed using Kruskal-Wallis tests with post-hoc Dunn’s tests, Wilcoxon matched pairs signed rank tests, or paired t tests as appropriate. A p-value < 0.05 was considered statistically significant. GraphPad Prism 10 was used for graphing and statistical analyses. Biorender.com was used for schematics prepared in Figures 1A and 2A. Sample sizes are specified in figure legends.

**Figure 1:**
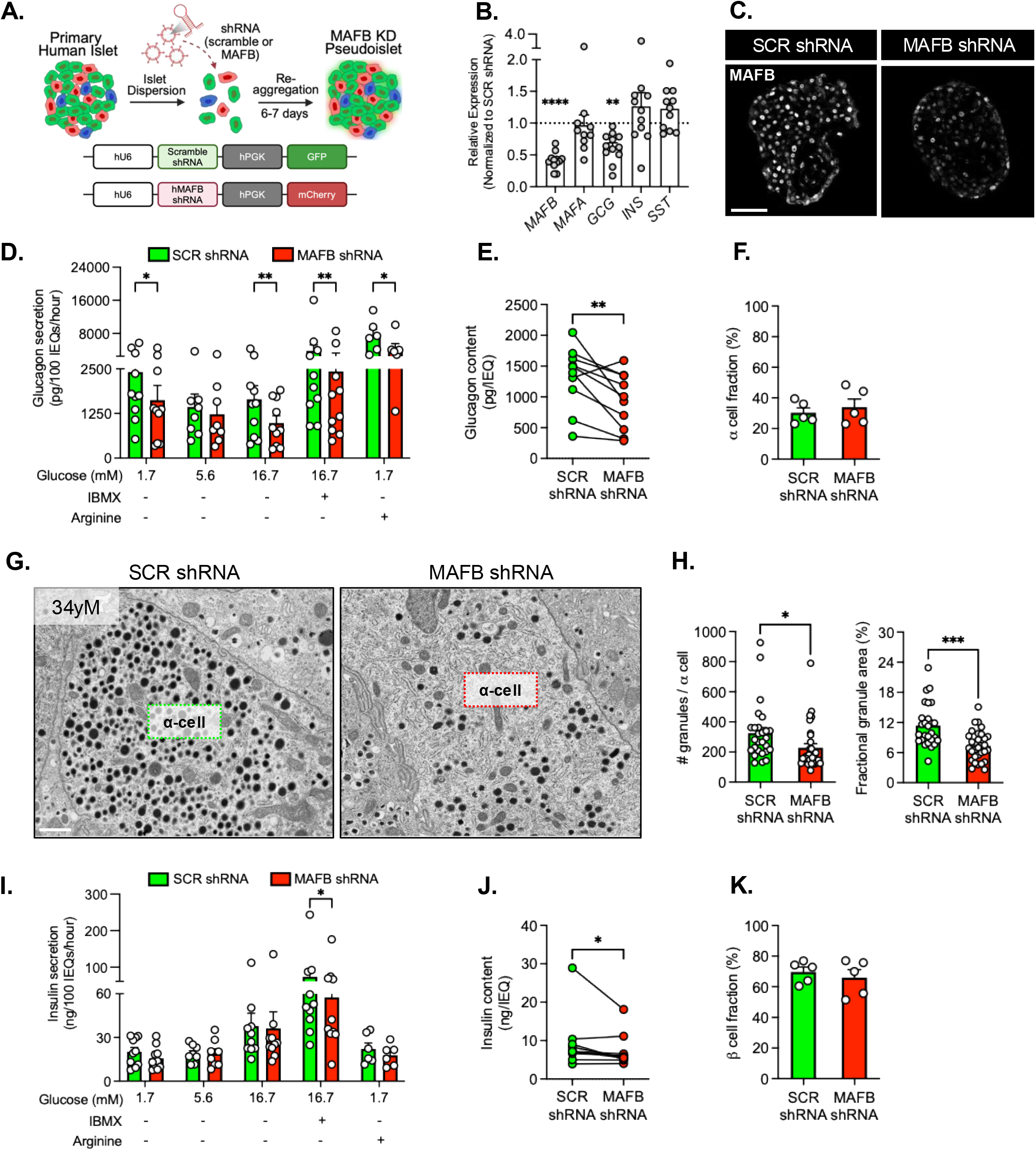
Acute MAFB knockdown reduces glucagon synthesis and secretion while only modestly compromising β-cell function. (A) Schematic of pseudoislet formation and lentiviral constructs coding for a Scramble (SCR) shRNA (nontargeting control) with a GFP reporter and a *MAFB* shRNA (MAFB knockdown [KD]) with an mCherry reporter. (B) Relative expression of *MAFB*, *MAFA*, and islet hormone genes in MAFB KD pseudoislets, quantified by qRT-PCR and normalized to SCR shRNA controls (dotted line; n = 10-12 donors per gene). (C) Representative immunofluorescent (IF) images of SCR shRNA (left) and MAFB shRNA (right) pseudoislets stained for MAFB. Scale bar: 50 µm. (D) Glucagon secretion by SCR shRNA (green bars) and MAFB shRNA (red bars) pseudoislets in static incubation experiments at low glucose (1.7 mM), basal glucose (5.6 mM), high glucose (16.7 mM), high glucose with cAMP stimulation (16.7 mM + 100 μM isobutylmethylxanthine [IBMX]), and low glucose with amino acid stimulation (1.7 mM glucose + 20 mM arginine) across 6-10 independent donors (2-3 technical replicates per condition per donor). Glucagon secretion was normalized to pseudoislet volume expressed in islet equivalents (IEQs). (E) Glucagon content normalized to IEQ in SCR shRNA (green circles) and MAFB shRNA (red circles) pseudoislets. (F) Average α-cell proportion as determined by IF analysis of whole pseudoislets stained for glucagon in SCR shRNA (green bar) and MAFB shRNA (red bar) groups (up to 10 pseudoislets per donor across 5 independent donors). (G) Representative TEM micrographs (2700x magnification) of a single α-cell in SCR shRNA (left) and MAFB shRNA (right) pseudoislets acquired from a 34-year-old male donor without diabetes. Scale bar: 1 µm. (H) Quantification of TEM images showing the average number (left) and fractional area (right) of glucagon-containing granules in SCR shRNA (green bars) and MAFB shRNA (red bars) α-cells (n = 27 α-cells and 8710 glucagon granules counted in SCR shRNA; n= 31 α-cells and 7082 glucagon granules counted in MAFB shRNA across 3 independent donors). (I) Insulin secretion by SCR shRNA (green bars) and MAFB shRNA (red bars) pseudoislets in static incubation experiments at low glucose (1.7 mM), basal glucose (5.6 mM), high glucose (16.7 mM), high glucose with cAMP stimulation (16.7 mM + 100 μM IBMX), and low glucose with amino acid stimulation (1.7 mM glucose + 20 mM arginine) across 6-10 independent donors (2-3 technical replicates per condition per donor). Insulin secretion was normalized to pseudoislet volume expressed in IEQs. (J) Insulin content normalized to IEQ in SCR shRNA (green circles) and MAFB shRNA (red circles) pseudoislets. (K) Average β-cell proportion as determined by IF analysis of whole pseudoislets stained for c-peptide in SCR shRNA (green bar) and MAFB shRNA (red bar) groups (up to 10 pseudoislets per donor across 5 independent donors). Data are presented as mean values ± SEM. One-way ANOVA (B), Wilcoxon matched pairs signed rank tests (D, E, H, I, J), and paired t tests (F, K) were used to generate *P* values: \**P* < 0.05, \*\**P* < 0.01, \*\*\**P* < 0.001, \*\*\*\**P* < 0.0001.

**Figure 2:**
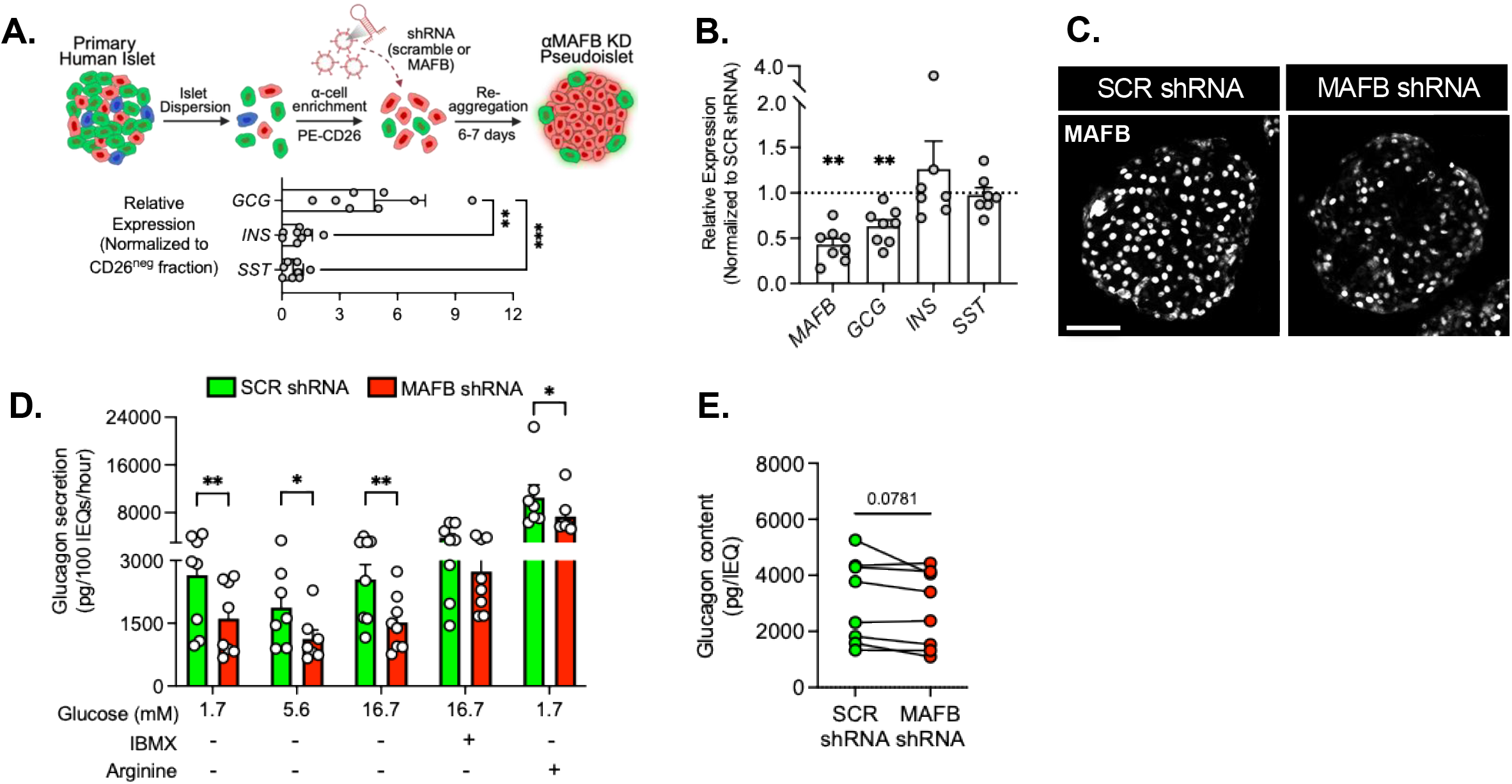
CD26^+^ α-cell enrichment uncovers MAFB-regulated defects in glucagon secretory capacity. (A) Schematic for α-cell-enriched pseudoislet formation and assessment of islet hormone gene expression in the CD26^+^ α-cell fraction by qRT-PCR (n = 8). (B) Relative expression of *MAFB*, *MAFA*, and islet hormone genes in MAFB KD pseudoislets, quantified by qRT-PCR and normalized to SCR shRNA controls (dotted line; n = 7-8 donors per gene). (C) Representative immunofluorescent (IF) images of SCR shRNA (left) and MAFB shRNA (right) α-cell-enriched pseudoislets stained for MAFB. Scale bar: 50 µm. (D) Glucagon secretion by SCR shRNA (green bars) and MAFB shRNA (red bars) α-cell-enriched pseudoislets in static incubation experiments at low glucose (1.7 mM), basal glucose (5.6 mM), high glucose (16.7 mM), high glucose with cAMP stimulation (16.7 mM + 100 μM IBMX), and low glucose with amino acid stimulation (1.7 mM glucose + 20 mM arginine) across 8 independent donors (2-3 technical replicates per condition per donor). Glucagon secretion was normalized to pseudoislet volume expressed in IEQs. (E) Glucagon content normalized to IEQ in SCR shRNA (green circles) and MAFB shRNA (red circles) α-cell-enriched pseudoislets. Data are presented as mean values ± SEM. Kruskal-Wallis test (A), one-way ANOVA (B), and Wilcoxon matched pairs signed rank tests (D, E) were used to generate *P* values: \**P* < 0.05, \*\**P* < 0.01, \*\*\**P* < 0.001.

### Data and resource availability

The RNA-Seq datasets will be available in the National Center for Biotechnology Information’s Gene Expression Omnibus (GEO) database). Further requests for resources and reagents should be directed to and will be fulfilled by the Lead Contact.

## RESULTS

### Glucagon secretion is disrupted upon MAFB KD in whole pseudoislets

MAFB is the most broadly and abundantly expressed large MAF TF in human pancreatic islets (Supp. Fig. 1a), correlates positively with electrophysiological activity markers in human α- and β-cell subpopulations^9^ (Supp. Fig. 1b) and is downregulated in islets from organ donors with diabetes.^5,7,8^ In contrast, the other large MAFs, NRL and C-MAF, are minimally expressed in the islet and MAFA is enriched in β-cells (Supp. Fig. 1a). To test the hypothesis that compromised MAFB activity elicits dysfunctional islet phenotypes, we used a pseudoislet approach^19,20^ to knockdown (KD) *MAFB* expression in primary human islets, thereby modeling the decrease seen in disease states. Following dispersion, primary islet cells were transduced with lentiviruses coexpressing a fluorescent transgene and short hairpin RNA (shRNA) directed against human *MAFB* (MAFB shRNA) or a scrambled control sequence (SCR shRNA), then reaggregated to form pseudoislets (Fig. 1a). Six days later, MAFB KD was found to reduce mRNA (i.e., by 59±4%) and protein levels which resulted in a 38±7% decrease in *GCG* expression but did not impact *MAFA, INS*, or *SST* mRNA production (Fig. 1b-c). Since MAFA shares cis-element target sequences with MAFB and is enriched in adult β-cells, the selective downregulation of *GCG* mRNA suggests that MAFA buffers β-cells against acute MAFB deficiency, whereas α-cells, which lack MAFA or other potential compensatory large MAF TFs, are more vulnerable to MAFB depletion.^15,46^

To assess the impact of MAFB depletion on α-cell function, we performed static incubation experiments and measured hormone secretion in SCR control and MAFB KD pseudoislets under basal (5.6 mM glucose) and regulated conditions (1.7 mM glucose, 16.7 mM glucose, 16.7 mM glucose + 100 μM IBMX, and 1.7 mM glucose + 20 mM arginine) (Fig. 1d). In SCR controls, glucagon secretion normalized to pseudoislet cell volume (expressed in islet equivalents [IEQs]) showed the expected glucose-dependent suppression, IBMX-mediated potentiation, and arginine stimulation (Fig. 1d). In contrast, MAFB-deficient pseudoislets exhibited lower glucagon secretion across all secretagogues, accompanied by a 30±8% reduction in hormone content. This reduction was not attributable to a decrease in the percentage of glucagon-positive α-cells as determined by immunohistochemical analysis but rather reflected reduced hormone stores per cell (Fig. 1e-f). Transmission electron microscopy corroborated these findings, revealing fewer glucagon granules and reduced fractional granule area in MAFB-deficient α-cells (Fig. 1g-h). Importantly, fractional glucagon secretion (normalized to glucagon content) did not differ between groups (Supp. Fig. 2a), indicating that the lower glucagon secretion in MAFB KD whole pseudoislets results from reduced hormone content.

### Whole pseudoislet MAFB KD only modestly impacts β-cell secretion

MAFB and MAFA coexpression marks the most functionally mature islet β-cell populations^9^, while MAFB knockout in human ES derived β-cells prevents insulin expression.^10^ To determine whether MAFB depletion impairs adult β-cell function in primary human islets, we measured insulin secretion in the static incubation experiments outlined above. When normalized to IEQ, insulin secretion was largely comparable between groups across most secretagogues, except for a modest reduction in IBMX-potentiated insulin secretion in MAFB KD pseudoislets (Fig. 1i). Fractional insulin secretion (normalized to insulin content) did not differ between groups under any condition (Supp. Fig. 2b). Insulin content was reduced by 16±6% on average (Fig. 1j); however, this difference was driven largely by a single donor, indicating that acute MAFB KD has a small and inconsistent effect on β-cell insulin stores. The fraction of islet β-cells also did not differ between groups (Fig. 1k). Collectively, these findings reveal that MAFB is essential for α-cell secretory activity and hormone content while having only minimal effects on adult β-cell function *in vitro*, most likely due to MAFA compensatory activity. We therefore focused our efforts on determining how MAFB regulates α-cells.

### CD26^+^ α-cell enrichment uncovers a cell-autonomous role for MAFB in glucagon secretion

Glucagon secretion in the intact islet is shaped by a dense network of autocrine, paracrine, and nutrient inputs that regulate α-cell excitability, cAMP signaling, and hormone biosynthesis through continuous crosstalk with β-, δ-, and non-endocrine neighbors.^47^ In this network context, whole pseudoislet MAFB KD cannot distinguish α-cell intrinsic effects from those arising through the surrounding islet environment. To resolve this, we applied a recently developed positive selection strategy that enriches for primary human α-cells.^26^ Dispersed human islets were labeled with a PE-conjugated antibody against CD26, an α-cell-selective surface protein^48,49^, separated from the CD26-negative (CD26^-^) fraction using magnetic beads, transduced with MAFB shRNA (αMAFB^KD^) or SCR shRNA (αSCR) lentiviruses, and reaggregated into pseudoislets (Fig. 2a). CD26-positive (CD26^+^) fractions showed ∼4-fold enrichment of *GCG*-expressing cells over CD26^-^ fractions (Fig. 2a) and a 5-fold increase in the glucagon:insulin ratio (Supp. Fig. 2c-d). Relative to αSCR controls, αMAFB^KD^ pseudoislets exhibited reductions in *MAFB* mRNA (57±7% decrease) and protein alongside a 37±7% decrease in *GCG* expression (Fig. 2b-c), confirming efficient and impactful MAFB depletion in α-cell-enriched pseudoislets.

Static incubation experiments revealed significantly lower glucagon secretion in αMAFB^KD^ pseudoislets across basal (5.6 mM), high (16.7 mM), and low (1.7 mM) glucose conditions, with or without arginine stimulation (Fig. 2d). Unlike whole pseudoislet MAFB KD, αMAFB^KD^ pseudoislets exhibited significantly impaired fractional glucagon secretion (Supp. Fig. 2e) and only a borderline reduction in glucagon content (P=0.07; Fig. 2e). Because this fractional deficit emerged when α-cells were removed from the broader islet milieu, it reveals a cell-autonomous role for MAFB in the α-cell secretory machinery, downstream of glucagon biosynthesis. Presumably this is due to the loss of paracrine inputs from neighboring islet cells, which antagonize glucagon production and secretion in whole pseudoislets.^47^

### Bulk RNA-seq of αMAFB^KD^ pseudoislets reveals compromised α-cell identity and lineage drift

To define the mechanism(s) underlying MAFB-regulated α-cell dysfunction, we performed bulk RNA-seq on these αMAFB^KD^ and αSCR control pseudoislets generated from adult human donors (Supp. Table 1). DESeq2^35^ identified 848 differentially expressed protein-coding genes (DEG; 414 down, 434 up; adjusted P < 0.05; Fig. 3a-b; Supp. Table 4). Gene set overlap analysis of the top 200 most significantly up- and down-regulated genes against the Human Molecular Signatures Database^36,37,50^ revealed that compromising MAFB levels disrupts α-cell identity while concurrently activating mesenchymal programs and inducing alternative islet endocrine fates (Fig. 3c-d). We use the term *lineage drift* to describe this coordinated loss of mature α-cell identity coupled with ectopic activation of mesenchymal and alternative endocrine programs, encompassing the related concepts of dedifferentiation and lineage infidelity. Downregulated genes reflected a marked loss of α-cell-enriched signatures, with nearly 20% mapping to pancreatic α-cell gene sets (Fig. 3c). These included canonical α-cell markers (*GCG, TTR, PCSK2, CRYBA2, LOXL4, FAP, CFC1*, *PLCE1*) and neuroendocrine secretory machinery spanning synaptic vesicle fusion (*SYT5, GAP43, ATP1A3, APLP1, PTPRN*), secretory vesicle components (*SYNGR4, SYT11, TRH*), and neurogenesis-associated factors (*CRTAC1, RAB29, THRB, NEFH, LIMK1*). Their distribution across synaptic, vesicular trafficking, and neuronal developmental processes indicates that MAFB maintains an integrated program coupling hormone production with the neuronal-like exocytotic machinery required for regulated glucagon secretion. MAFB also regulated genes involved in amino acid transport (*SLC7A8, SLC38A4, SLC3A2*) and small molecule metabolism (*PAPSS2, PRPS1, PDHB, GPD1, TAT*) (Fig. 3c). Notably, 25 downregulated genes are Polycomb SUZ12 targets bearing repressive H3K27me3 epigenetic marks (e.g., *PCSK2, CRTAC1, CRYBA2, NEFH, LBH*). This suggests that MAFB antagonizes PRC2-mediated methyltransferase activity that is normally involved in controlling expression of genes critical to α-cell differentiation.

**Figure 3:**
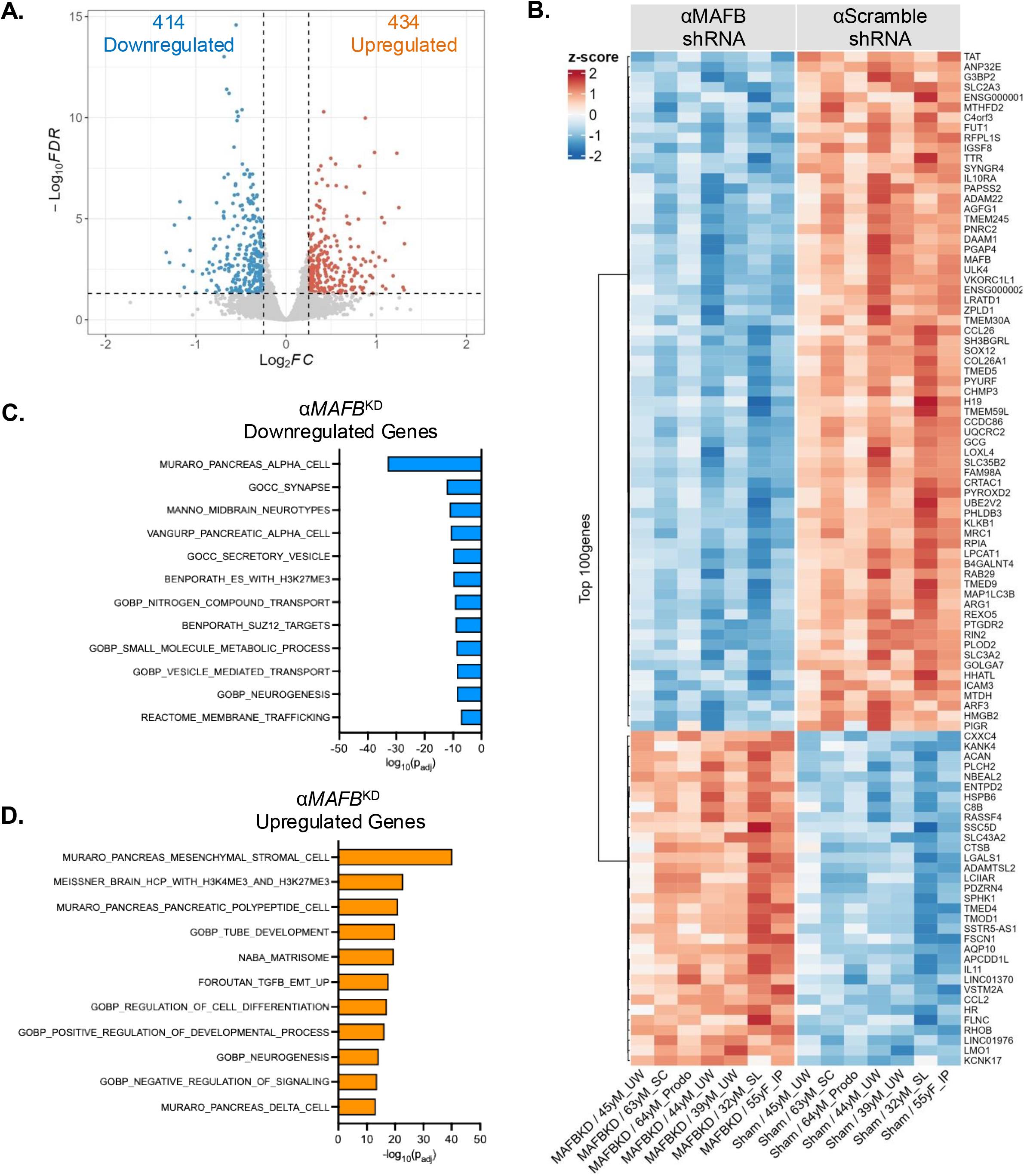
Bulk transcriptomic profiling of CD26^+^ *MAFB*-deficient pseudoislets reveals compromised identity and lineage drift in primary human α-cells. (A) Volcano plot of DEGs in αMAFB^KD^ pseudoislets. (B) Heat map of the top 100 most significant DEGs in αMAFB^KD^ versus αSCR control pseudoislets. Columns represent individual donor samples; rows represent genes. Bar graphs showing significantly downregulated (C) and upregulated (D) gene set overlap enrichment terms in αMAFB^KD^ pseudoislets, queried against the Human MSigDB database using hypergeometric testing with Benjamini-Hochberg FDR correction (q < 0.05).

### MAFB KD activates mesenchymal and alternative endocrine programs

In rodent β-cells, loss of mature identity TFs (e.g., FoxO1^51^ and Pdx1^52^) triggers dedifferentiation toward progenitor-like states and acquisition of α-cell features, and human T2D α- and β-cells similarly lose mature identity and derepress juvenile programs.^53,54^ Strikingly, MAFB KD in αMAFB^KD^ islets simultaneously activated mesenchymal/stromal signatures and upregulated genes associated with alternative islet endocrine cell types (Fig. 3d). The most enriched upregulated gene sets included mesenchymal/stromal markers (*ITGA5, MXRA8, MRC2, PRRX1*), extracellular matrix (ECM) and matrix-remodeling genes (*COL6A2*, *LOX, LOXL2, TGFBI, HAS2*), epithelial-mesenchymal transition (EMT) components (*VIM, SPHK1*), and TGF-β/Wnt ligands implicated in fibrogenesis and developmental plasticity (WNT5A, INHBA) alongside Notch target and vascular/endothelial-associated genes (*JAM3, HEYL*). In addition, overlap analysis revealed enrichment for pancreatic polypeptide (PP-cell) and δ-cell signatures, indicating that MAFB deficiency permits ectopic expression of alternative endocrine programs.

Prior work shows that MAFB loss drives a shift toward δ- and PP-cell identity at the expense of β- and α-cells^10^ in MAFB-depleted hESC-derived endocrine cells and produces a marked increase in PP-cells in pancreatic endocrine-specific MafB KO mice.^55^ Our findings extend this pattern to adult primary human α-cells, supporting a conserved role for MAFB in restricting alternative endocrine fates across developmental and adult contexts. This δ-/PP-directed reprogramming is phenotypically distinct from that caused by loss of the α-cell specification factor ARX, which in adult mouse α-cells drives conversion toward β-cell fates^56^, identifying MAFB and ARX as nonredundant α-cell maintenance factors that guard against distinct alternative identities.

The broad representation of mesenchymal, ECM-remodeling, and developmental genes across cell migration and tissue development gene sets suggests that MAFB normally suppresses mesenchymal transition and tissue remodeling programs. Consistent with this, 39 upregulated genes overlapped with a bivalently marked (H3K4me3/H3K27me3) chromatin domain gene set (MSigDB MEISSNER_BRAIN_HCP_WITH_H3K4ME3_AND_H3K27ME3). Overlapping genes included developmental regulators (*TBX3, GLI2*), endocrine lineage TFs (*ISL1, MAF*), and mesenchymal genes, implying that MAFB prevents derepression of plasticity genes residing in poised chromatin domains (Fig. 3d). Collectively, these data establish MAFB as a critical transcriptional regulator that maintains α-cell identity by coordinating activation of lineage-specific genes with suppression of mesenchymal and alternative endocrine programs.

### No significant MAFB-dependent changes were detected by scRNA-seq of adult human β-cells

To resolve the bulk transcriptional changes at cellular resolution and to test whether MAFB KD also alters β-cell transcriptomes, we performed scRNA-seq on matched SCR control and MAFB KD whole pseudoislets from two healthy donors (Supp. Table 1), yielding 1,491 high-quality β-cell transcriptomes and 4,464 high-quality α-cell transcriptomes (Fig. 4a; Supp. Fig. 3a-b). Notably, MAFB KD produced no significant DEGs in β-cells other than *MAFB* itself, indicating that acute MAFB deficiency does not profoundly remodel the β-cell transcriptome under these conditions. This molecular observation parallels the modest insulin secretion phenotype in the whole pseudoislet MAFB KD group.

**Figure 4:**
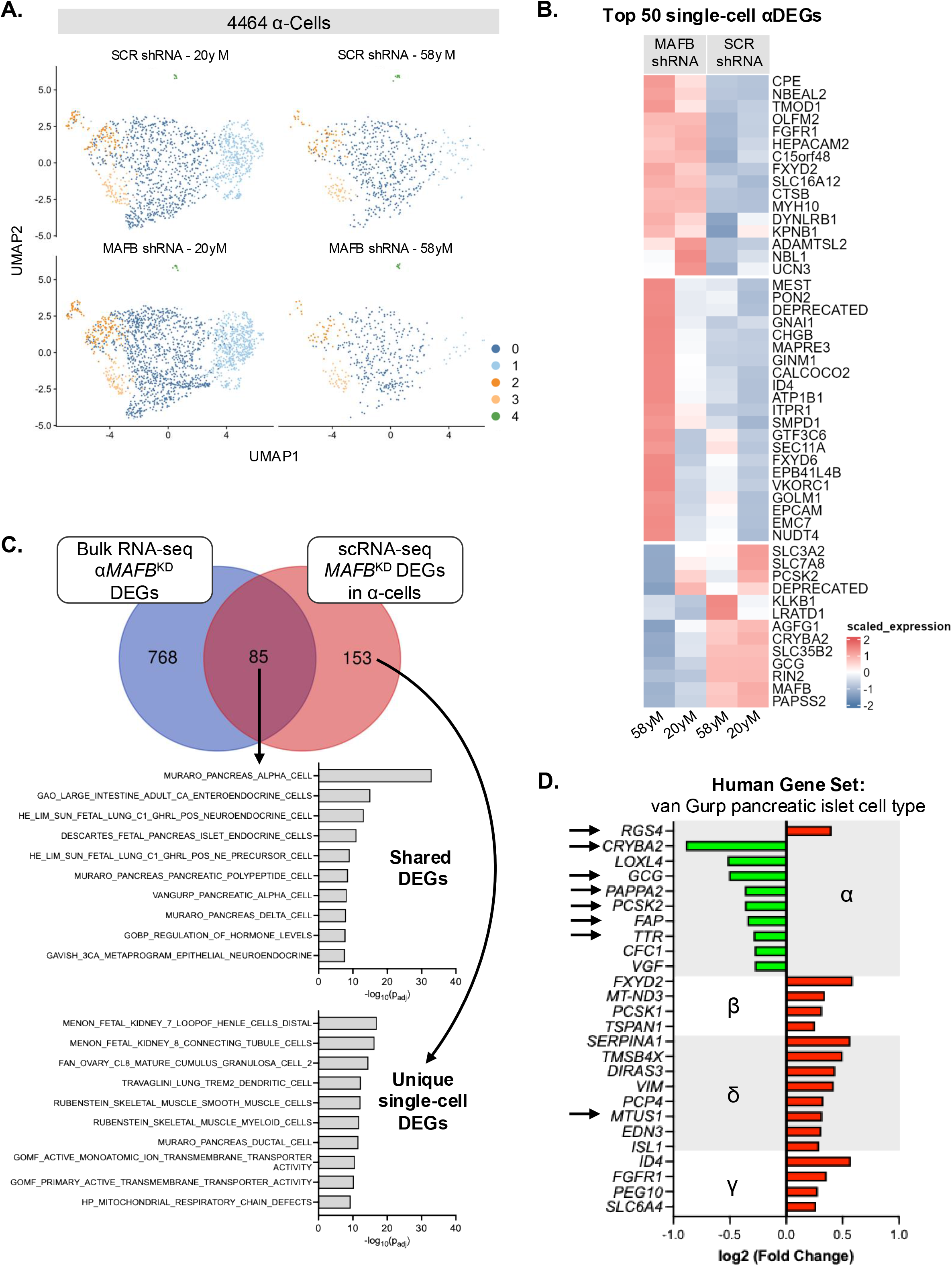
MAFB KD drives α-cell transcriptional reprogramming toward alternative islet cell fates. (A) UMAP projections of 4,464 α-cells from SCR shRNA and MAFB KD whole pseudoislets across 2 independent donors (20yM and 58yM). Cells are colored by subcluster identity (α-clusters 0-4). Total α-cells per sample: SCR shRNA 20yM, n = 669; MAFB KD 20yM, n = 429; SCR shRNA 58yM, n = 1,430; MAFB KD 58yM, n = 1,936. (B) Heat map of scaled expression for the top 50 most significant α-cell DEGs between MAFB KD and SCR shRNA conditions. Columns represent individual donor samples; rows represent genes. (C) Venn diagram depicting the overlap in MAFB-regulated α-cell DEGs between α-cell-enriched bulk (αMAFB^KD^ DEGs) and scRNA-seq datasets. 763 DEGs were uniquely identified in bulk αMAFB^KD^ pseudoislets, 153 DEGs were uniquely identified at single-cell resolution, and 85 DEGs were shared between datasets. Bar graphs show gene set overlap enrichment terms for shared DEGs (top) and unique single-cell DEGs (bottom), queried against the Human MSigDB database using hypergeometric testing with Benjamini-Hochberg FDR correction (q < 0.05). (D) Log_2_ fold change of significant DEGs from the van Gurp human islet cell type-specific genesets^59^ in MAFB KD versus SCR shRNA α-cells. Genes are grouped by islet cell type. Green bars indicate downregulation; red bars indicate upregulation. Arrows denote putative direct transcriptional targets of MAFB, as defined by previously published ChIP-seq data from primary human islets.^60,102^

### MAFB maintains α-cell identity across the α-cell compartment

In contrast to β-cells, scRNA-seq analysis of α-cells identified 238 protein-coding DEGs in MAFB KD versus SCR controls (49 down, 189 up; adjusted P < 0.05; Fig. 4b; Supp. Table 5). These overlapped substantially but incompletely with our bulk α-cell-enriched dataset: 85 genes (∼36%) were shared, while 153 (∼64%) were unique to single-cell resolution (Fig. 4c). The smaller DEG count and partial overlap likely reflect the well-documented limitations of droplet-based scRNA-seq and the smaller donor n^57,58^, such that the two analyses offer complementary views of α-cell remodeling.

The 85 shared DEGs showed striking coherence, with marked overrepresentation of pancreatic α-cell identity signatures (31 genes) spanning canonical markers and secretory machinery (*GCG, PCSK2, TTR, CFC1, PAPSS2, CRYBA2, FAP, LOXL4*, *SNAP25, SYT4, APLP1, CHGB*) and related endocrine signatures (Fig. 4c [shared]). Consistent with these aggregate changes, profiling of islet cell-type-specific identity genes in individual α-cells confirmed that MAFB KD downregulated canonical α-cell-signature genes^59^ (*CRYBA2, LOXL4, GCG, PAPSS2, PCSK2, FAP, TTR, CFC1, VGF*), many of them putative direct MAFB targets (arrows) based on whole islet ChIP-seq^60^, while inducing ectopic expression of genes normally restricted to β- (*FXYD2, MT-ND3, PCSK1, TSPAN1*), δ- (*SERPINA1, TMSB4X, DIRAS3, VIM, PCP4, MTUS1, EDN3, ISL1*), and γ-cells (*ID4, FGFR1, PEG10, SLC6A4*) (Fig. 4d). These data corroborate, at single-cell resolution, that MAFB acts cell-autonomously to maintain α-cell identity by activating α-specific genes and repressing alternative endocrine programs.

Notably, the 153 unique single-cell DEGs defined regulatory pathways absent from the bulk analysis, including kidney tubule, pancreatic ductal, β-cell, and ion transporter signatures (Fig. 4c [unique]), suggesting that MAFB deficiency also remodels transcriptional programs that vary across α-cell subsets and are masked when α-cells are analyzed in aggregate. We therefore examined MAFB-dependent gene regulation at the level of individual α-cell subpopulations.

### MAFB orchestrates α-cell subpopulation-specific changes

Human α-cells are functionally and molecularly heterogeneous^9,54,61^, with cells co-expressing MAFB and ARX predicted to be the most functionally mature relative to those expressing only one or neither factor.^9^ To resolve how MAFB shapes this heterogeneity, we performed differential expression and pathway analysis within each of the five single-cell α-cell clusters in SCR control and MAFB KD (C0-C4; Fig. 4a; Supp. Table 6). Cluster C4 (33 α-cells) yielded no significant DEGs, focusing downstream analyses on C0-C3 (Fig. 5a-b; Supp. Fig. 3c). This revealed both common and cluster-restricted mechanisms of MAFB-dependent control.

**Figure 5:**
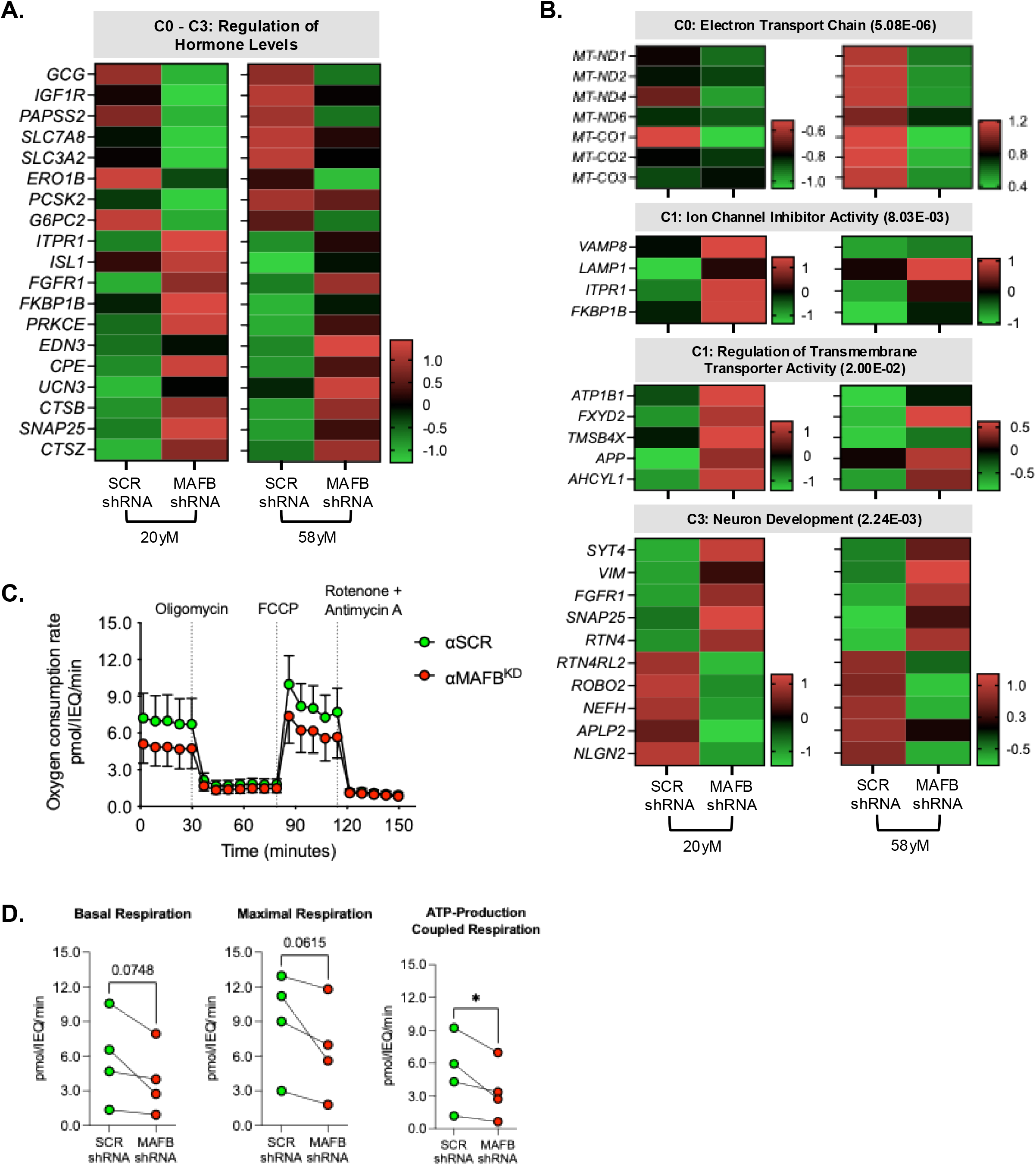
MAFB regulates subcluster-specific transcriptional programs in MAFB KD α-cells, including a mitochondrial respiration signature validated by pseudoislet respirometry. (A) Heatmap of scaled expression for DEGs enriched in the shared g:Profiler pathway term “Regulation of Hormone Levels” across α-cell subclusters C0-C3, comparing SCR shRNA and MAFB KD conditions in two donors (20yM and 58yM). (B) Heatmaps of scaled expression for representative α-cell subcluster-specific DEGs enriched in top g:Profiler pathway terms, identified by independent per-subcluster DE analysis: Electron Transport Chain (C0; adjusted *P* = 5.08 x 10^-6^), Ion Channel Inhibitor Activity (C1; adjusted *P* = 8.03 x 10^-3^), and Neuron Development (C3; adjusted *P* = 2.24 x 10^-3^). Adjusted *P*-values shown in panel headers. Color scales indicate z-scored normalized expression. (C) Seahorse XF Pro mitochondrial stress test trace of oxygen consumption rate (OCR; pmol/IEQ/min) in α-cell-enriched SCR shRNA (green circles) and MAFB KD (red circles) pseudoislets. Dotted vertical lines indicate sequential injection of oligomycin, FCCP, and rotenone/antimycin A. (D) Quantification of basal respiration (left), maximal respiration (middle), and ATP-production coupled respiration (right) derived from the Seahorse assay in (C), comparing SCR shRNA and MAFB KD α-cell-enriched pseudoislets (n = 4 independent donors). Each point represents an individual donor; lines connect paired observations. Data are presented as mean values ± SEM. *P* values: Statistical significance was determined by ratio paired t-test. \**P* < 0.05; *P*-values shown for trending comparisons.

All four clusters (C0-C3) shared enrichment of the “Regulation of Hormone Levels” pathway upon compromising MAFB levels, indicating that MAFB controls core metabolic and secretory functions across α-cell populations (Fig. 5a). Common downregulated genes included those required for metabolic sensing and glucagon secretion (*GCG, IGF1R, PAPSS2, SLC7A8, SLC3A2, ERO1B, PCSK2, G6PC2*), whereas upregulated genes comprised calcium signaling regulators (*ITPR1, FKBP1B, PRKCE*), neuropeptides (*EDN3, UCN3*), synaptic proteins (*SNAP25*), and developmental lineage factors (*FGFR1, ISL1*).

Three clusters (C0, C1, and C3) additionally displayed cluster-restricted MAFB-regulated pathways undetectable at bulk resolution (Fig. 5b). In the larger C0 population, MAFB KD reduced expression of mitochondrially encoded electron transport chain (ETC) subunits (*MT-ND1, MT-ND2, MT-ND4, MT-ND6, MT-CO1, MT-CO2, MT-CO3*), consistent with a role in maintaining mitochondrial oxidative metabolism in this large α-cell subset. In C1, MAFB KD increased expression of regulators of the autophagic-lysosomal machinery (*VAMP8, LAMP1*), intracellular calcium-handling (*ITPR1, FKBP1B*), ion homeostasis and the cytoskeleton (*ATP1B1, FXYD2, TMSB4X, APP, AHCYL1*), implicating MAFB in vesicular trafficking and calcium dynamics relevant to stimulus-secretion coupling. In C3, MAFB KD increased neuronal and secretory plasticity markers (*SYT4, VIM, FGFR1, SNAP25, RTN4*) and decreased genes that regulate mature neuronal architecture and synaptic organization (*RTN4RL2, ROBO2, NEFH, APLP2, NLGN2*), pointing to MAFB-linked maintenance of a mature neuroendocrine state. These findings suggest that MAFB operates as a transcriptional orchestrator combining uniform control of core hormonal programs across all α-cells with cluster-restricted programs that preserve specialized functional competence.

### MAFB maintains mitochondrial respiration in adult human α-cells

To test whether the MAFB-associated downregulation of mitochondrially encoded ETC genes in the C0 population produces a functional respiratory defect, we measured oxygen consumption rates in αSCR and αMAFB^KD^ pseudoislets. MAFB KD significantly reduced ATP-linked respiration in α-cell-enriched pseudoislets, with concordant trends toward reduced basal (P=0.07) and maximal (P=0.06) respiration (Fig. 5c-d). A parallel defect was evident in whole MAFB KD pseudoislets, which showed reduced basal and ATP-linked respiration versus SCR controls (Supp. Fig. 4a-b). Together, these data link MAFB-dependent ETC gene expression to mitochondrial respiratory capacity in adult human α-cells and indicate that the bioenergetic consequences of MAFB deficiency extend to the intact islet.

## DISCUSSION

MAFB is downregulated in T1D and T2D α- and β-cells and is enriched in the most functionally mature islet cell subpopulations^5–9^, but its role in adult human α- and β-cell function has not been directly tested. This gap reflects a broader limitation in the field: islet-enriched TFs have been characterized predominantly in rodent developmental and conditional deletion models, which leave species differences unresolved, and in human stem cell-derived islets, which capture endocrine cell fate specification rather than mature cell maintenance.^62^ The distinction between establishing and sustaining islet cell identity may be particularly consequential for MAFB, whose expression pattern diverges between species and persists into adulthood in human α- and β-cells.^9,10,13–16^ Here, we combined shRNA-mediated MAFB KD in whole- and α-cell-enriched human pseudoislets with functional and molecular assays to address this gap. Acute MAFB KD produced only limited functional changes in adult human β-cells, whereas the same perturbation destabilized α-cell identity, impaired glucagon synthesis and secretion, and compromised mitochondrial respiration. This asymmetry reframes how MAFB decline likely contributes to α-versus β-cell dysfunction in diabetes.

### Acute MAFB KD produces limited β-cell changes in whole pseudoislets

β-cell responses to MAFB KD were far more modest than those of α-cells. Insulin content was slightly reduced and IBMX-potentiated insulin release slightly attenuated, but fractional insulin secretion and β-cell fraction were preserved, and scRNA-seq identified no additional β-cell DEGs beyond *MAFB* itself. Somatostatin RNA levels in MAFB expressing δ-cells were likewise unaffected (Fig. 1b). This α-versus β-cell asymmetry is the inverse of that reported during human stem cell differentiation, where MAFB is indispensable for β-cell derivation but only partially required for α-cells.^10^ One plausible explanation is functional compensation by MAFA, which is enriched in β-cells but largely absent from α-cells and binds shared cis-elements at β-cell targets including *INS*.^17,63^ *MAFA* mRNA was unchanged in MAFB KD pseudoislets (Fig. 1b), suggesting that any such buffering would not require transcriptional upregulation of MAFA. Other explanations, including partial rather than complete MAFB depletion or the short (6-day) perturbation window, could equally account for the limited β-cell phenotype we observed. Notably, Cataldo et al.^64^ previously showed that combined MAFA/MAFB silencing in primary human islets and the human EndoC-βH1 cell line impairs insulin secretion and exocytotic gene expression, suggesting that large MAF activity is required in β-cells even if MAFB alone is not acutely rate-limiting. Whether chronic MAFB loss, more complete depletion, or some other perturbation would ultimately erode adult δ- and/or β-cell function remains an important open question.

### Acute MAFB KD preferentially disrupts α-cell glucagon regulation

A key finding of this study is that acute MAFB depletion preferentially disrupts glucagon regulation in primary human α-cells. In whole pseudoislets, MAFB KD reduced glucagon content by ∼30%, decreased both glucagon granule number and fractional area, and suppressed total glucagon secretion across all secretagogues tested, while fractional glucagon secretion was preserved. This pattern suggests that MAFB primarily supports glucagon biosynthesis, with impaired total secretion largely reflecting reduced cellular glucagon stores rather than a proportional defect in secretory efficiency. Consistent with this interpretation, islet glucagon content is the strongest predictor of glucagon secretion in isolated human islets, in contrast to insulin.^65^ These findings also align with prior rodent studies showing reduced glucagon content and impaired glucose-regulated glucagon secretion following MafB loss in adult α-cells.^14,15,55,66^ Mechanistically, MafB has been shown to directly activate the glucagon gene through binding to the conserved G1 cis-acting element in the rodent αTC-6 cell line.^15^ Together, out data extend this conserved regulatory relationship to adult human α-cells, establishing MAFB as a conserved regulator of glucagon production.

### CD26^+^ enrichment reveals α-cell-autonomous secretory defects

To disentangle α-cell autonomous MAFB requirements from those transmitted through the intact islet, we turned to the CD26^+^ α-cell-enriched pseudoislet system.^26^ This approach uncovered a dimension of MAFB regulation not apparent in whole islet experiments. In CD26^+^ αMAFB^KD^ pseudoislets, fractional glucagon secretion was significantly impaired, revealing that MAFB maintains not only glucagon biosynthesis but also the secretory machinery required for stimulus-secretion coupling. The relatively preserved fractional glucagon secretion in whole pseudoislets likely reflects regulation by non-α endocrine cells and/or paracrine signaling.^26^ Upon α-cell enrichment, this buffering is reduced, unmasking a cell-autonomous requirement for MAFB in α-cell secretory capacity. This α-cell-autonomous secretory defect is especially relevant to islet dysfunction in T1D and T2D, contexts in which the secretion of key paracrine regulators from neighboring endocrine cells (e.g. insulin from β- and somatostatin from δ-cells), is disrupted, further limiting the buffering that partially compensates for α-cell defects in the intact islet. ^67–72^ Interestingly, this pattern parallels recent findings for RFX6, another α-cell-enriched TF that is dysregulated in diabetic human islets.^26,30^ Coykendall et al.^26^ showed that RFX6 KD restricted to CD26^+^ α-cells impaired exocytosis and glucagon secretion, whereas these phenotypes were absent after whole islet RFX6 suppression. That the same whole islet versus α-cell-targeted discordance emerges for two independent α-cell TFs argues that cell-type-selective perturbation is essential for dissecting TF function in heterogeneous human islets, and points to paracrine reinforcement of α-cell identity within the intact islet.^70^

### MAFB maintains α-cell identity and suppresses lineage plasticity

Bulk transcriptomic profiling of CD26^+^ αMAFB^KD^ pseudoislets revealed that MAFB sustains an integrated transcriptional program linking α-cell identity to secretory competence. Nearly 20% of all downregulated genes mapped to pancreatic α-cell gene sets, spanning canonical markers (*GCG, TTR, PCSK2, CRYBA2*), neuroendocrine secretory machinery (*SYT5, GAP43, PTPRN*), and amino acid transporters (*SLC7A8, SLC38A4*). This coordinated loss indicates that MAFB does not merely activate individual α-cell genes but maintains the functional architecture coupling hormone production, regulated exocytosis, and metabolic sensing.

Equally striking was the concurrent activation of mesenchymal and alternative endocrine lineage programs in MAFB-deficient α-cell-enriched pseudoislets, including ECM remodeling genes (*COL6A2, LOX, LOXL2*), EMT-associated components (*VIM, SPHK1*), and δ- and PP-cell markers. This composite signature is reminiscent of, though not identical to, the dedifferentiation paradigm developed primarily in rodent β-cells^51^, in which loss of lineage-defining TFs permits re-emergence of developmentally silenced and alternative endocrine programs. Evidence for analogous processes in human islets is more limited and context-dependent: T2D donor pancreata show hormone-negative endocrine cells and mislocalized FOXO1/NKX6.1^73^, T2D α-cells de-repress an immature, juvenile transcriptional program with impaired exocytosis^53,54^, and T1D α-cells show reduced ARX and MAFB, ectopic β-cell TF expression (i.e., PDX1 or NKX6.1) in a fraction of GCG^+^ cells, and rare bihormonal GCG^+^/INS^+^ cells.^8,56^ The EMT component of our signature parallels β-cell findings in which miR-7-mediated dedifferentiation engages a Pdx1-Ovol2-Zeb2 axis to drive EMT-associated gene expression and fibrotic remodeling^74^, and with the observation by Avrahami et al.^53^ that adult human α-cells retain immature features even in donors without diabetes. In T2D, both α- and β-cells undergo further maturity loss, de-repressing juvenile and exocrine/mesenchymal programs alongside stress- and inflammation-response signatures to yield an immature, mixed-lineage transcriptome.^53,54,75^ Together, our findings support a model in which adult α-cell identity is actively maintained by MAFB, whose loss unmasks latent developmental and mesenchymal programs.

### MAFB KD engages a mesenchymal drift signature

The identity destabilization we observe in MAFB KD α-cells closely parallels the recently proposed mesenchymal drift framework^76^, which defines age- and disease-associated identity loss as a directionally biased accumulation of mesenchymal programs in hybrid cell states that retain lineage features, with identity-defining TFs serving as barriers against drift. Our α-cell lineage drift signature aligns with each pillar of this framework. First, MAFB KD α-cells enriched for TGF-β, Wnt, Notch, and matrisome programs, the core signaling and structural signatures of mesenchymal drift. Second, mesenchymal markers were acquired in α-cells that retained glucagon and core α-cell identity, consistent with a hybrid intermediate rather than complete lineage conversion. Third, mitochondrially encoded ETC subunits were coordinately downregulated with impaired ATP-linked respiration, paralleling the oxidative-to-glycolytic shift central to mesenchymal drift. Fourth, upregulated genes were enriched at bivalent H3K4me3/H3K27me3 loci and PRC2 targets were enriched among downregulated genes, consistent with the epigenetic remodeling that accompanies drift. Notably, the epigenetic architecture of α-cells may underlie this vulnerability. Bramswig et al.^77^ previously showed that human α-cells carry substantially more bivalent marks than β-cells, with histone methyltransferase inhibition inducing glucagon/insulin co-expression in human islets. Our findings suggest that MAFB normally represses these bivalent developmental and plasticity loci, reinforcing α-cell identity at the epigenomic level and providing a mechanistic candidate for why α-cells drift toward mesenchymal and alternative endocrine fates when MAFB is depleted.

These observations situate MAFB within a broader paradigm in which islet endocrine cell fate stability requires continuous transcriptional reinforcement. In β-cells, conditional TF deletion of Pdx1^52^, Nkx6.1^78,79^, Pax6^80,81^, Isl-1^82^, or Nkx2.2^83^ causes loss of β-cell identity and ectopic activation of alternative endocrine programs. The corresponding α-cell network (e.g., Arx^56,84,85^, Pax6^86–90^, and Nkx2.2/Klf4^91^) has been less well defined. Our findings extend this paradigm to MAFB and to the adult human α-cell, and we predict that complete and chronic MAFB depletion will amplify the identity destabilization reported here, potentially advancing adult α-cells beyond the hybrid intermediate toward more stable lineage conversion.

### Single-cell analysis reveals subpopulation specific MAFB regulation and a MAFB mitochondrial axis

Single-cell transcriptomics revealed that MAFB regulates both shared and subpopulation specific programs across heterogeneous α-cell clusters. Beyond corroborating our bulk findings, this analysis uncovered 153 DEGs visible only at single-cell resolution, enriched for ion transporter, ductal, and kidney signatures, thereby exposing regulatory effects masked in aggregate populations. Distinct cluster-level responses emerged in the α-cell compartment: cluster 0 selectively downregulated mitochondrial ETC genes; cluster 1 remodeled vesicular trafficking and calcium signaling; and cluster 3 activated neuronal differentiation programs. These findings suggest that MAFB deficiency acts non-uniformly, preferentially compromising specific α-cell subsets and offering a plausible transcriptional framework for the heterogeneous α-cell dysfunction observed in T2D.

Of these cluster-restricted programs, the ETC signature was the most directly testable, and cluster 0 was also the largest α-cell subpopulation. We therefore asked whether this transcriptional change was accompanied by a measurable respiratory deficit. Indeed, MAFB KD significantly impaired ATP-linked respiration in α-cell-enriched pseudoislets, with concordant trends toward reduced basal and maximal respiration. These data establish that MAFB is required to sustain mitochondrial respiratory capacity in adult human α-cells.This MAFB-mitochondrial relationship is echoed across disease-relevant human datasets. MAFB is reduced in T1D and T2D α-cells^5,8^; single-cell studies show coordinated MAFB and ETC dysregulation in T2D α-cells^7^; patch-seq links mitochondrial respiratory chain complex assembly to α-cell exocytotic capacity in health and its disruption in T2D^54^; and ER stress in stem-cell-derived α-cells uniquely downregulates MAFB with coordinated glycolytic and oxidative phosphorylation suppression.^4^ That acute MAFB KD alone reproduces features of these signatures suggests that reduced MAFB may be sufficient to initiate, rather than merely accompany, α-cell bioenergetic decline.

The correspondence across datasets is not uniform, however. Whereas the MAFB downregulation reported by Bosi et al.^7^ aligns with the ETC-low state of our cluster 0, Dai et al.^54^ also identified a dysfunctional T2D α-cell subset with paradoxically elevated MAFB and other lineage markers, raising the possibility that MAFB elevation in some disease contexts marks failed functional maturation. Whether our ETC-low cluster 0 represents an early transcriptional state preceding the electrophysiological dysfunction captured by patch-seq, or a molecularly distinct subpopulation, remains to be determined. Even so, the convergence of these datasets positions the MAFB-ETC axis as a candidate link between α-cell transcriptional dysregulation and the bioenergetic failure characteristic of diabetic islets.

Although we did not test whether reduced respiration is required for the secretory phenotypes we observe, the two are plausibly linked. Regulated glucagon secretion is energetically demanding, requiring ATP for the metabolic signaling that couples nutrient sensing to α-cell excitability, for granule biogenesis and prohormone processing, for the priming of secretory granules that precedes calcium-triggered fusion, and for the ion pumps that maintain the electrochemical and calcium gradients on which stimulus-secretion coupling depends.^92–98^ Notably, α-cell glucagon release depends on ATP in a bell-shaped rather than a linear manner, being suppressed at both insufficient and excessive ATP levels.^96,98^ We speculate that a MAFB-dependent reduction in oxidative capacity narrows the energetic reserve available to support the transcriptional loss of secretory machinery and prohormone processing genes found upon MAFB KD.

In summary, this study identifies MAFB as a cell-autonomous regulator of adult human α-cell maintenance, coordinating glucagon biosynthesis, stimulus-secretion coupling, identity preservation, lineage stability, and mitochondrial function. Acute MAFB KD was far more disruptive to α-cells than β-cells, indicating that diabetes-associated MAFB downregulation may be especially consequential for α-cell dysfunction. By linking MAFB-dependent α-cell identity programs to subcluster-specific ETC gene expression and bioenergetic competence, these findings position MAFB as a non-redundant node in the adult human α-cell transcriptional network and provide a framework for understanding how α-cell dysfunction occurs in T1D and T2D islets.

### Limitations of the study

Several limitations should be considered. First, our shRNA-based KD approach achieves partial rather than complete MAFB depletion. While this models the graded loss observed in disease, it may underestimate the full scope of MAFB-dependent regulation. Second, the 6-day pseudoislet culture period captures short-term transcriptional and functional consequences but may not recapitulate the chronic, progressive nature of MAFB loss in diabetes. Likewise, our functional studies were performed under non-stressed conditions and do not address whether MAFB-dependent pathways are differentially engaged under the metabolic stress conditions characteristic of the diabetic milieu. Lastly, our scRNA-seq analysis was performed on two donors, which limits detection of more subtle transcriptional changes.

## ETHICS DECLARATIONS

## Acknowledgements

We gratefully acknowledge the organ donors and their families for their invaluable gift, which made this research possible. Human pancreatic islets and/or other resources were provided by the NIDDK-funded Integrated Islet Distribution Program (IIDP) (RRID:SCR _014387) at City of Hope, NIH Grant # U24DK098085 and the JDRF-funded IIDP Islet Award Initiative. This manuscript used RNA-seq data obtained from PanKbase^99,100^ (https://www.pankbase.org; accessed 20251010), supported by NIDDK U24DK138515, U24DK138512, and supplemental funds from the NIH Office of Data Science Strategies. This dataset represents isolated pancreatic islet tissue from human samples from the Human Pancreas Analysis Program^22–25^ (HPAP; RRID:SCR_016202), IIDP, and Prodo. HPAP is part of a Human Islet Research Network (HIRN; RRID:SCR_014393) with funding from UC4-DK112217, U01-DK123594, UC4-DK112232, and U01-DK123716. This manuscript also includes data from HumanIslets.com^101^ funded by the Canadian Institutes of Health Research, JDRF Canada, and Diabetes Canada (5-SRA-2021-1149-S-B/TG 179092) with data from islets isolated by the Alberta Diabetes Institute IsletCore at the University of Alberta in Edmonton (www.bcell.org/adi-isletcore) with the assistance of the Human Organ Procurement and Exchange (HOPE) program, Trillium Gift of Life Network (TGLN), and other Canadian organ procurement organizations. All donors’ families gave informed consent for the use of pancreatic tissue in research as approved by the Human Research Ethics Board at the University of Alberta (Pro00013094). Transmission electron microscopy was performed using the Vanderbilt Cell Imaging Shared Resource (supported by NIH grants CA68485, DK20593, DK58404, DK59637, EY08126, 1S10OD034315-01, and R1R24OD037694-01). We sincerely thank the Vanderbilt Creative Data Solutions Shared Resource (RRID:SCR_022366) team, especially Matthew Cottam, Shristi Shrestha, and Jean-Philippe Cartailler, for performing bulk and scRNA-seq data processing and analysis. The Agilent Seahorse XF Pro Analyzer is housed and managed within the Vanderbilt High-Throughput Screening Core Facility, an institutionally supported core, and was funded by NIH Shared Instrumentation Grant 1S10OD018015. Vanderbilt Technologies for Advanced Genomics (VANTAGE) core laboratory is supported in part by Clinical and Translational Science Award Grant 5UL1 RR024975-03, Vanderbilt Ingram Cancer Center Grant P30 CA68485, Vanderbilt Vision Center Grant P30 EY08126, and National Institutes of Health/National Center for Re-search Resources Grant G20 RR030956.

## Data availability

The RNA-Seq datasets will be available in the National Center for Biotechnology Information’s Gene Expression Omnibus (GEO) database). All other data associated with this manuscript can be found in the manuscript, in the ESM files.

## Contribution statement

KCC and RWS conceived and designed the study. Funding was obtained by KCC, RWS, ACP, SKK, and PEM. KCC, JL, MG, XT, CH, ND, GR, NH, REJ, and RA performed experiments and analyzed the data. JPC performed computational analyses. KCC and RWS interpreted the data and wrote the manuscript. JPC, VMNC, ACP, PEM, and SKK made contributions to the conception or design of the work, provided materials, and/or edited the manuscript. All authors read and approved the final manuscript. KCC and RWS serve as guarantors of this work, and both have verified the underlying data of this manuscript.

---

**Supplemental Figure 1:**
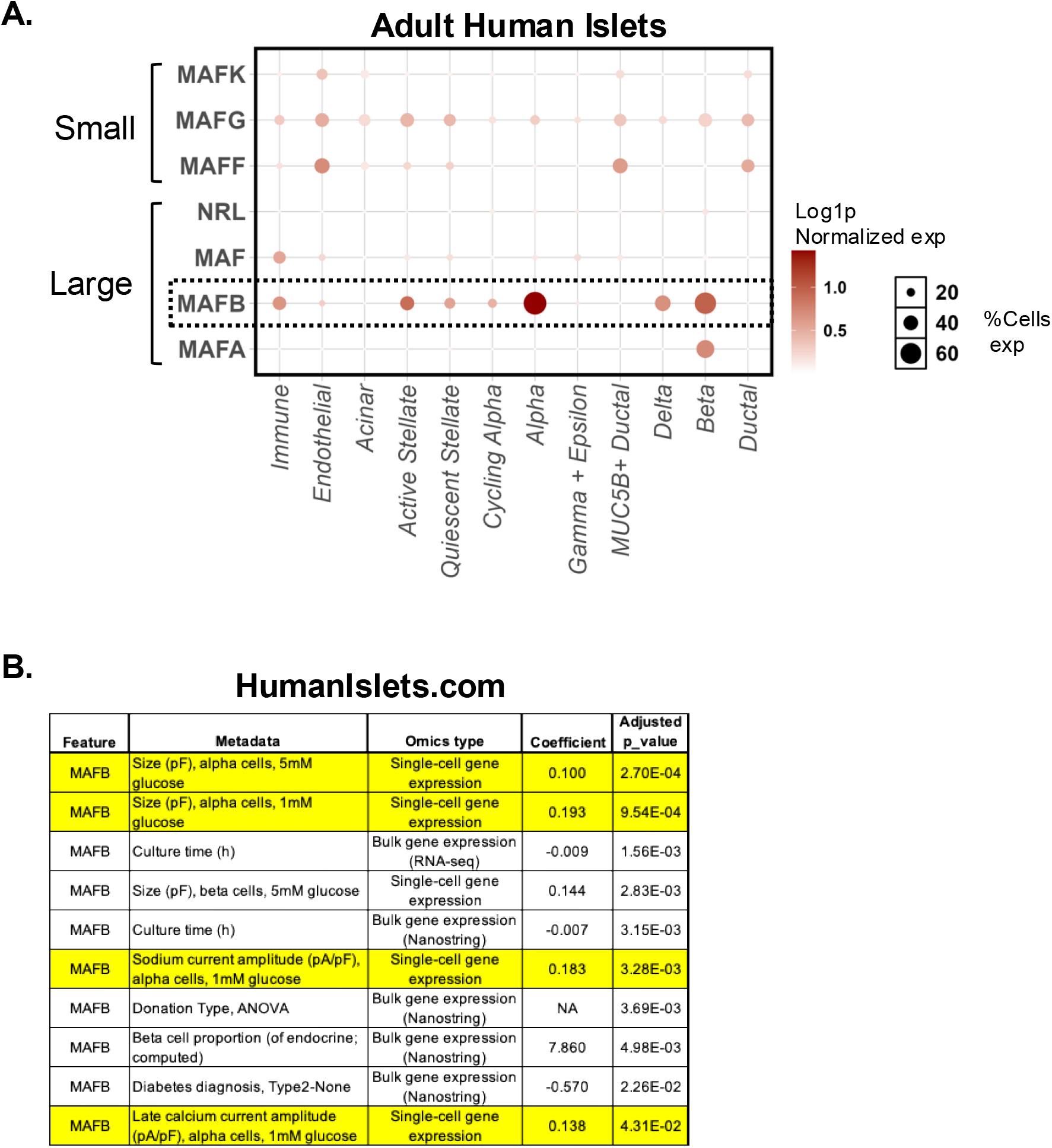
MAFB is the predominant large Maf transcription factor in human islets and correlates with α- and β-cell electrophysiological activity. (A) Dot plot of single-cell-resolved normalized gene expression of the Maf family of transcriptional regulators in isolated human pancreatic islets from PanKbase^99,100^ (https://www.pankbase.org; accessed 20251010). Small, small Maf proteins; Large, large Maf proteins. Dotted line shows *MAFB* expression across all cell types, highlighting clear enrichment in α-cells. (B) Table highlighting features positively correlated with α-cell MAFB expression in isolated human islets from HumanIslets.com.^101^

**Supplemental Figure 2:**
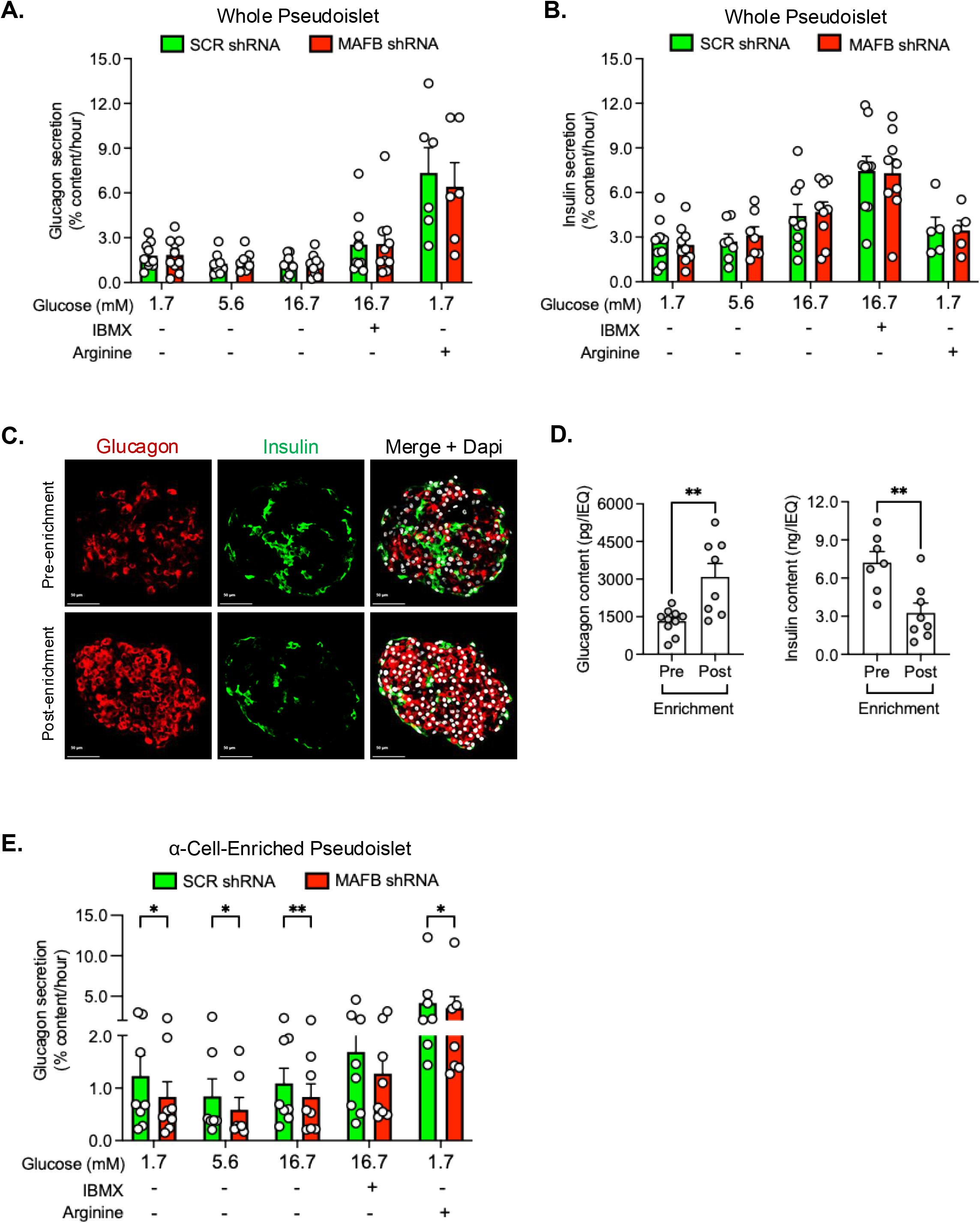
Islet α-cell enrichment unmasks a fractional glucagon secretory impairment upon MAFB deficiency. Fractional glucagon (A) and insulin (B) secretion by SCR shRNA (green bars) and MAFB shRNA (red bars) whole pseudoislets in static incubation experiments at low glucose (1.7 mM), basal glucose (5.6 mM), high glucose (16.7 mM), high glucose with cAMP stimulation (16.7 mM + 100 μM IBMX), and low glucose with amino acid stimulation (1.7 mM glucose + 20 mM arginine) across 6-10 independent donors (2-3 technical replicates per condition per donor). Hormone secretion was normalized to glucagon (A) or insulin (B) content. (C) Representative IF images of a human pseudoislet stained for glucagon (red), insulin (green) and Dapi (white) pre- (top row) and post- (bottom row) CD26-positive α-cell enrichment. Scale bars: 50 µm. (D) Comparison of IEQ-normalized glucagon (left) and insulin (right) content pre- and post- CD26-positive α-cell enrichment in human pseudoislets across 8 independent donors. (E) Fractional glucagon secretion by SCR shRNA (green bars) and MAFB shRNA (red bars) α-cell-enriched pseudoislets in static incubation experiments at low glucose (1.7 mM), basal glucose (5.6 mM), high glucose (16.7 mM), high glucose with cAMP stimulation (16.7 mM + 100 μM IBMX), and low glucose with amino acid stimulation (1.7 mM glucose + 20 mM arginine) across 8 independent donors (2-3 technical replicates per condition per donor). Glucagon secretion was normalized to glucagon content. Data are presented as mean values ± SEM. Paired t test (D) and Wilcoxon matched pairs signed rank tests (A, B, E) were used to generate *P* values: \**P* < 0.05, \*\**P* < 0.01.

**Supplemental Figure 3:**
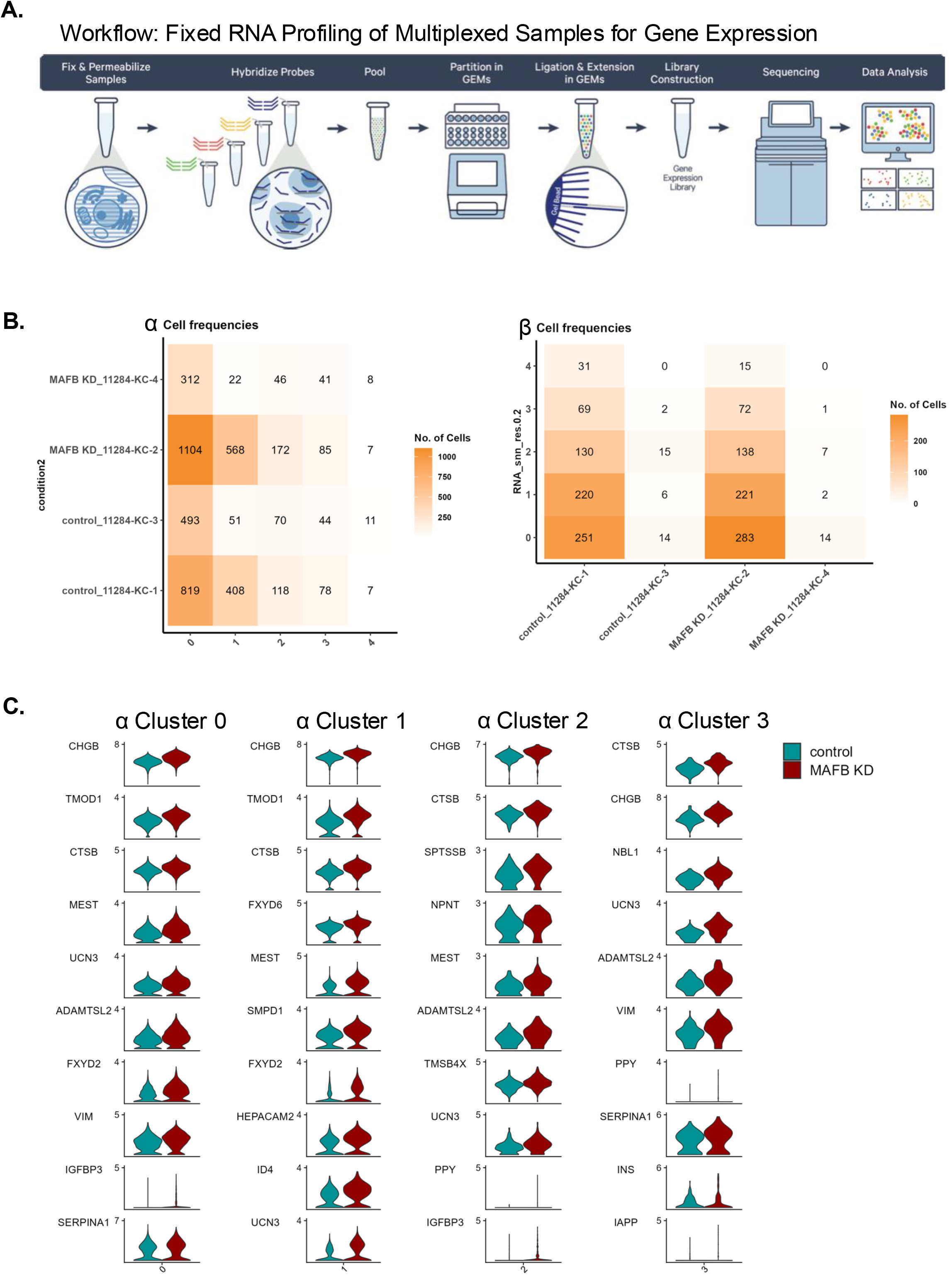
Single-cell fixed RNA profiling workflow and cluster-resolved gene expression in MAFB-deficient human α-cells. (A) Schematic of the 10x Genomics Fixed RNA Profiling workflow for multiplexed single-cell gene expression analysis. Control and MAFB KD human pseudoislets were fixed and permeabilized, hybridized with probe sets, pooled, and partitioned into gel beads-in-emulsion for ligation, extension, library construction, sequencing, and downstream data analysis. (B) Heatmaps showing the distribution of α- (left) and β- (right) cell frequencies across transcriptionally defined α clusters (0-4; left) or across each sample (right). (C) Violin plots showing expression of the top 10 DEGs in α-cell subclusters C0-C3 between SCR control (teal) and MAFB KD (burgundy) α-cells.

**Supplemental Figure 4:**
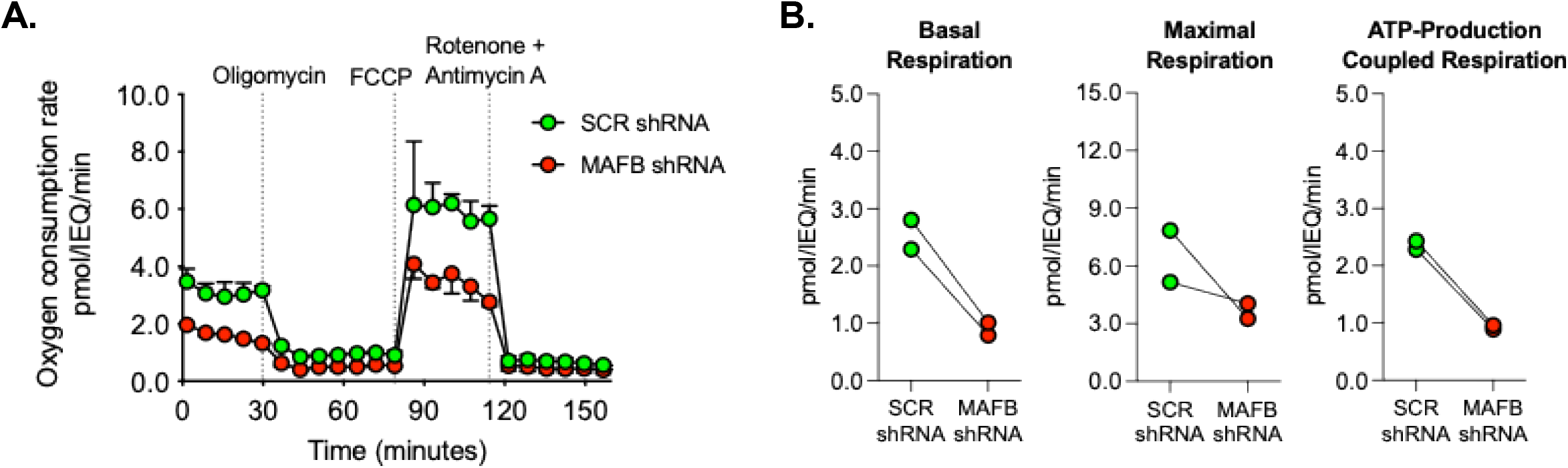
MAFB KD reduces mitochondrial respiration in whole pseudoislets. (A) Seahorse XF Pro mitochondrial stress test trace showing oxygen consumption rate (OCR; pmol/IEQ/min) over time in whole pseudoislets transduced with SCR shRNA (green circles) or MAFB-targeting shRNA (red circles). Dotted lines indicate sequential injections of oligomycin, FCCP, and rotenone + antimycin A to assess distinct components of mitochondrial respiration. (B) Quantification of basal respiration (left), maximal respiration (middle), and ATP-production coupled respiration (right) derived from the OCR traces in (A), comparing SCR shRNA (green circles) and MAFB KD (red circles) pseudoislets from n = 2 independent donor preparations. Each point represents an individual donor; lines connect paired observations. Data are presented as mean values ± SEM.

